# Transcription termination promotes splicing efficiency and fidelity in a compact genome

**DOI:** 10.1101/2025.03.12.642901

**Authors:** Keaton Barr, Kevin L He, Andreas J. Krumbein, Guillaume F. Chanfreau

## Abstract

Splicing of terminal introns is coupled to 3′-end processing by cleavage and polyadenylation (CPA) of mRNAs in mammalian genes. Whether this functional coupling is universally conserved across eukaryotes is unclear. Here we show using long read RNA sequencing in *S*.*cerevisiae* that splicing inactivation does not result in widespread CPA impairment. We also show that inactivation of CPA has limited impact on splicing efficiency. The negative impact of CPA inactivation on splicing is mainly due to transcription termination defects that promote readthrough transcription, leading to splicing inhibition for downstream intron-containing genes. Splicing impairment due to 5′ extensions is length-dependent and can be detected independently from CPA inactivation for endogenous or synthetic genes, and is likely due to an increased distance of splicing signals to the 5′ cap. Finally, we found that deficient termination can promote novel intragenic and long-range intergenic splicing events. These results argue against a broad coupling between splicing and CPA in *S*.*cerevisiae* but show that efficient CPA-mediated transcription termination is critical for splicing fidelity and efficiency in a compact genome.

**Significance Statement:** Accurate gene expression requires that the enzyme that polymerizes RNA stops at the proper site (termination). In addition multiple RNA processing reactions, including removal of intervening sequences are necessary to produce mature mRNAs. How these different steps in the RNA biogenesis pathways influence each other is not well understood. In this study, we show that inactivation of termination induces mature RNA formation defects, including long RNAs that retain intervening sequences, or chimeric RNAs containing sequences from genes located next to each other on the genome. This study underscores the importance of proper termination to ensure accurate and efficient splicing of adjacent genes, which is particularly critical for compact genomes in which genes are located close to each other.

## Introduction

Synthesis of most eukaryotic mRNAs requires three main processing steps: capping of the 5′-end of the mRNA by addition of a 7-methyl guanosine(m^7^G), removal of intronic sequences by splicing, and 3′-end formation by cleavage and polyadenylation (CPA). In addition to generating mRNA 3′-ends, CPA also promotes transcription termination by RNA polymerase II. The three major mRNA processing reactions occur co-transcriptionally, as factors involved are known to associate with RNA polymerase II and/or the nascent transcripts (reviewed in (1)). These three main processing reactions also influence each other. At the 5′-end of the mRNA, capping stimulates splicing of the first, cap-proximal intron ((2–4); reviewed in (5)). This stimulation is conferred by the cap-binding complex (CBC), which binds the m^7^G cap and promotes recruitment of the U1 snRNP, resulting in enhanced recognition of the first, cap-proximal exon 5′ splice sites (SS) by splicing factors (2, 3, 6).. At the 3′-end of the mRNA, splicing of the terminal intron and recognition of the CPA site are intimately connected (reviewed in (1, 7)). Interestingly, this coupling of splicing and 3′-end processing can occur independently from transcription(8, 9), and several interactions have been described between 3′-end processing factors and splicing factors. At the genomic scale, perturbation of 3′-end processing has been shown to result in inhibition of splicing of terminal introns(10). In addition, recent studies using long-read RNA sequencing have revealed a coordination between alternative splicing and polyadenylation in *Drosophila* (11), providing further evidence for the coupling between splicing and polyadenylation in metazoans genes.

While there is broad evidence for reciprocal stimulation of splicing of terminal introns and 3′-end processing by CPA in metazoans, it is unclear whether this mechanism is universally conserved across eukaryotes. In support of this conservation, inactivation of the *S*.*pombe* homologue of splicing factor U2AF65 triggers 3′-end formation defects and transcriptional read-through (12). In the yeast *S*.*cerevisiae*, most intron-containing genes contain only a single intron, which defines most introns as *de facto* ′terminal′ and close to the cleavage and polyadenylation sites, especially for introns of ribosomal protein genes which have small 3′-exons and constitute the quantitative majority of splicing substrates (13). Indirect evidence suggests that there might exist functional coupling between splicing and 3′-end formation in *S*.*cerevisiae*. The Ysh1 endonuclease involved in the cleavage reaction of CPA was first identified in a genetic screen for splicing mutants(14),(15). In addition, physical and genetic interactions have been identified between splicing factors and the CPA machinery (reviewed in (16)). Furthermore 3′-end processing and transcription termination defects induced by inactivation of the nuclear poly(A)-binding protein Nab2 resulted in an increase in intron retention(17).This observation suggested that defects in CPA-mediated transcription termination may impair intron recognition and promote splicing defects. However, no study has directly addressed the impact of inactivating CPA factors on splicing, or of inactivating splicing factors on 3′-end formation in *S*.*cerevisiae*. In this study we address this question using rapid inactivation of splicing or 3′-end processing factors followed by long-read RNA sequencing. We do not find evidence for a major impact of inactivating splicing on 3′-end processing. However, loss of termination caused by CPA inactivation results in intrusive transcription into downstream intron-containing genes, which decreased their splicing efficiency, as previously reported for Nab2 inactivation (17). This deleterious impact of long 5′-extensions on splicing was also observed independently from inactivating the CPA machinery for endogenous mRNAs with long 5′-extensions or for synthetic constructs. Furthermore, loss of termination revealed dormant splice sites and resulted in the activation of novel splice sites and the production of intergenic spliced mRNAs. These results underscore the importance of proper CPA-mediated termination to limit aberrant splicing events and enhance splicing efficiency in a genome where transcription units are located close to each other.

## Results

### Inactivation of the U1 snRNP component Snp1 does not lead to broad 3′-end formation or termination defects

To analyze the reciprocal impact of splicing or 3′-end processing factors on splicing and polyadenylation, we assessed the impact of inactivating these factors on the transcriptome. Most splicing or 3′-end processing factors are essential in *S*.*cerevisiae*. To rapidly inactivate these RNA processing factors, we used the anchor away (AA) technique(18), which promotes rapid export of a tagged nuclear protein out of the nucleus, resulting in functional inactivation. This technique has previously been used to inactivate splicing(19) or 3′-end processing (20, 21) factors in *S*.*cerevisiae*. We used AA strains of the 3′-end processing factors Ysh1 and Rna15, as well as the U1 snRNP component Snp1p, the yeast orthologue of the U1-70K protein. We chose to inactivate this early splicing factor to prevent engagement of transcripts in the splicing pathway and binding of proteins that may recognize splicing signals following E complex formation. Following treatment with rapamycin to induce the export of tagged proteins out of the nucleus, RNA processing efficiency was analyzed *in vivo* by long-read RNA sequencing. An untagged strain from the same genetic background (which alleviates any toxic effects of rapamycin treatment and downstream effects of gene expression) was treated for 2hrs with rapamycin and used as a negative control (mentioned as WT in the remainder of this study). RNA samples were obtained for three replicates for each strain and sequenced by direct RNA sequencing using the Oxford Nanopore Technology (ONT) platform. We obtained an average of ca. 3 million reads per replicate (Table S1) and excellent reproducibility of RNA detection between replicates (FigS1). Wes then analyzed the impact of inactivation of these factors on 3′-end formation by measuring changes in median 3′ UTR length (measured by the distance of the poly(A) site to the stop codon; see methods). Analysis of 3′-UTR lengths did not reveal any global impact of Snp1 inactivation (Fig1A,1B1). This is in contrast to Ysh1 or Rna15 inactivation, which resulted in a global increase in 3′-UTR size (Fig1C,1D), as expected since loss of these factors is known to induce termination defects. We note that a limited number of mRNAs showed a significant increase in median 3′ UTR length upon Snp1 inactivation (Fig1A,Table S2). Most of these genes exhibit multiple poly(A) sites in a wild-type context and we found that the increase in median 3′ UTR length was due to an increase in usage of distal poly(A) sites in the *snp1*-AA strain (See FigS2 for 2 examples, *COX6* and *RPB2*). However, the pattern of poly(A) site usage detected in the *snp1-AA* strain was overall more similar to the WT control than to the *ysh1*-AA or *rna15-AA* strains (FigS2), showing that the increase is UTR length is not due to an intrinsic defect in termination but rather to a subtle shift in poly(A) site usage. Since this increase in distal poly(A) site usage is limited to only a few mRNAs, we did not investigate further whether it is due to general splicing perturbation, or to a more specific role of Snp1. Importantly, none of the genes that showed an increase in median 3′ UTR length in the Snp1-AA strain contained an intron and, we did not detect any significant changes in 3′-UTR length for intron-containing genes (ICG) upon Snp1 inactivation (Fig1B). Based on these results we conclude that splicing inactivation at an early stage of the splicing pathway does not result in major perturbations of 3′-end processing *in vivo*.

**Figure 1.**
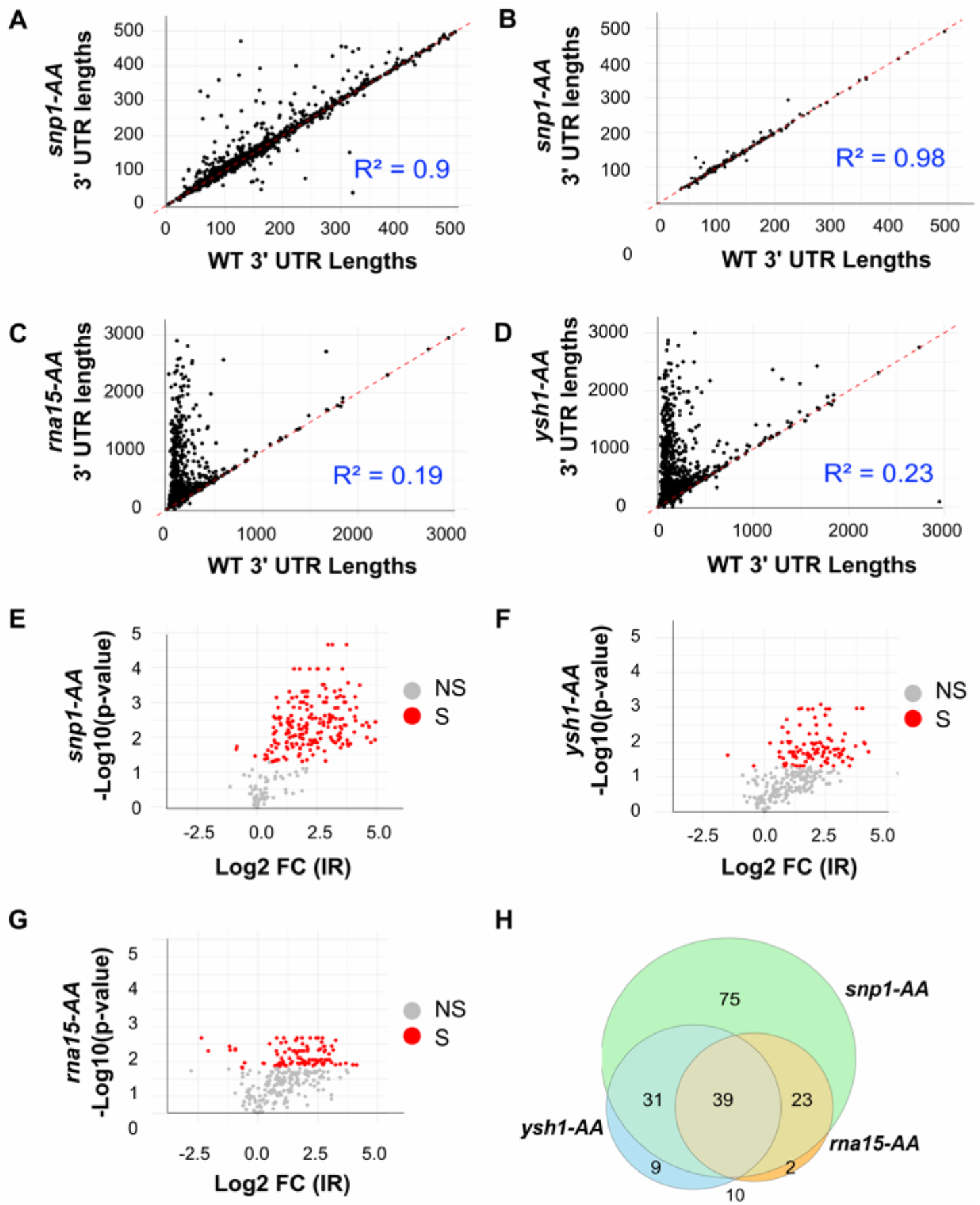
Global analysis of 3′-end formation and splicing upon CPA or splicing inactivation. A. Comparison of median 3′UTR length between the Snp1-AA strain and the WT control for all *S*.*cerevisiae* genes detected by Nanopore sequencing. Numbers indicate the average of the median 3’ UTR length for all 3 replicates. B. Comparison of median 3′UTR length between the *snp1*-AA strain and WT for all *S*.*cerevisiae* intron-containing genes. C. Comparison of median 3′UTR length between the *rna15*-AA and WT for all *S*.*cerevisiae* genes detected by Nanopore sequencing. D. Same as (C) for the *snp1*-AA strain. E. Volcano plot showing the fold-change in intron retention in the Snp1-AA strain compared to WT. Red dots correspond to ICGs showing significant splicing changes. F. Same as (E) for the *ysh1*-AA strain. G. Same as (E) for the *rna15*-AA strain. H. Venn Diagram showing the overlap of ICGs showing significant splicing defects in the *snp1*-AA, *snp1*-AA and *rna15*-AA strains.

### Inactivation of the cleavage and polyadenylation factors Ysh1 and Rna15 has limited impact on splicing efficiency

We next focused on analyzing the impact of inactivating Snp1, Ysh1 or Rna15 on pre-mRNA splicing. To this end, we quantitated unspliced reads in strains in which these factors were anchored away. Analysis of intron retention (IR) in the Snp1-AA strain (measured by counting the number of unspliced reads over the total number of reads for each ICG) showed an increase in IR for many ICGs (168), as expected (Fig1E). Some ICGs did not show significant accumulation of unspliced reads: either because their splicing is robust enough to resist to short-term inactivation of the U1 snRNP, or because unspliced pre-mRNAs accumulation is limited by rapid degradation (22, 23). By contrast, inactivation of Rna15 or Ysh1 had a more limited effect on intron retention (Fig1F, 1G, 1H), with 81 and 74 ICGs showing significant splicing defects in *ysh1*-AA and *rna15-* AA, respectively (Fig1H). Some splicing defects overlap was observed upon inactivation of Rna15 or Ysh1(Fig1H). However, inactivation of Ysh1 and Rna15 also resulted in a unique set of genes for which splicing was affected (Fig1H). This result suggests that each of these factors impact expression of the affected genes in a unique way which perturbs splicing indirectly, rather than through a general coupling mechanism in which splicing efficiency requires a direct recognition of nearly polyadenylation sites. We next focused on investigating the mechanisms by which splicing is perturbed upon inactivation of these CPA factors.

### Splicing defects are not prevalent in 3′-extended RNAs but can be masked by nuclear degradation

To understand the mechanistic basis for the perturbation of splicing upon Ysh1 or Rna15 inactivation, we first analyzed the splicing signals of ICGs showing a significant splicing defect in the *ysh1*-AA or *rna15*-AA samples. Sequence logos obtained for the 5′-splice site (SS) or branchpoint sequences of the RNAs with splicing defects in the *ysh1*-AA or *rna15*-AA strain were similar to those of all ICG (FigS3), suggesting that deficient splicing for these mRNAs was not due to intrinsically weaker splicing signals. To better understand the mechanism responsible for splicing inhibition, we inspected ONT reads of ICGs to search for the presence of unspliced RNAs in transcripts with 3′-readthroughs in the *ysh1*-AA or *rna15*-AA samples, as 3′-readthrough transcripts were shown previously to be more prone to splicing defects when transcription termination efficiency is reduced upon Nab2 nuclear depletion (17). In general, intron-containing transcripts with long 3′-extensions following Ysh1 or Rna15 inactivation did not show increased intron retention. One example is shown in Fig2A for *RPL26A*, which exhibits frequent 3′-readthrough transcripts onto the downstream gene *YLR345W*, none of which were unspliced. We calculated the fraction of intron retention for RNAs with 3′-extensions in the *ysh1*-AA or *rna15*-AA or control strain genome-wide and compared it to RNAs with no extensions. RNAs with 3′-extensions found in the *ysh1*-AA or *rna15*-AA datasets did not show a large increase of intron retention compared to unextended transcripts found in the control wild-type sample, or compared to unextended transcripts detected in the *ysh1*-AA or *rna15*-AA samples (Fig2B). Some exceptions include *RPL28*, for which intron retention was twice as abundant in 3′-extended transcripts compared to unextended transcripts in the *ysh1*-AA or *rna15*-AA (FigS4). Overall, these results suggest that loss of recognition of the primary CPA or termination site next to the intron has limited impact the efficiency of splicing of introns located just upstream of these poly(A) sites genome-wide.

**Figure 2.**
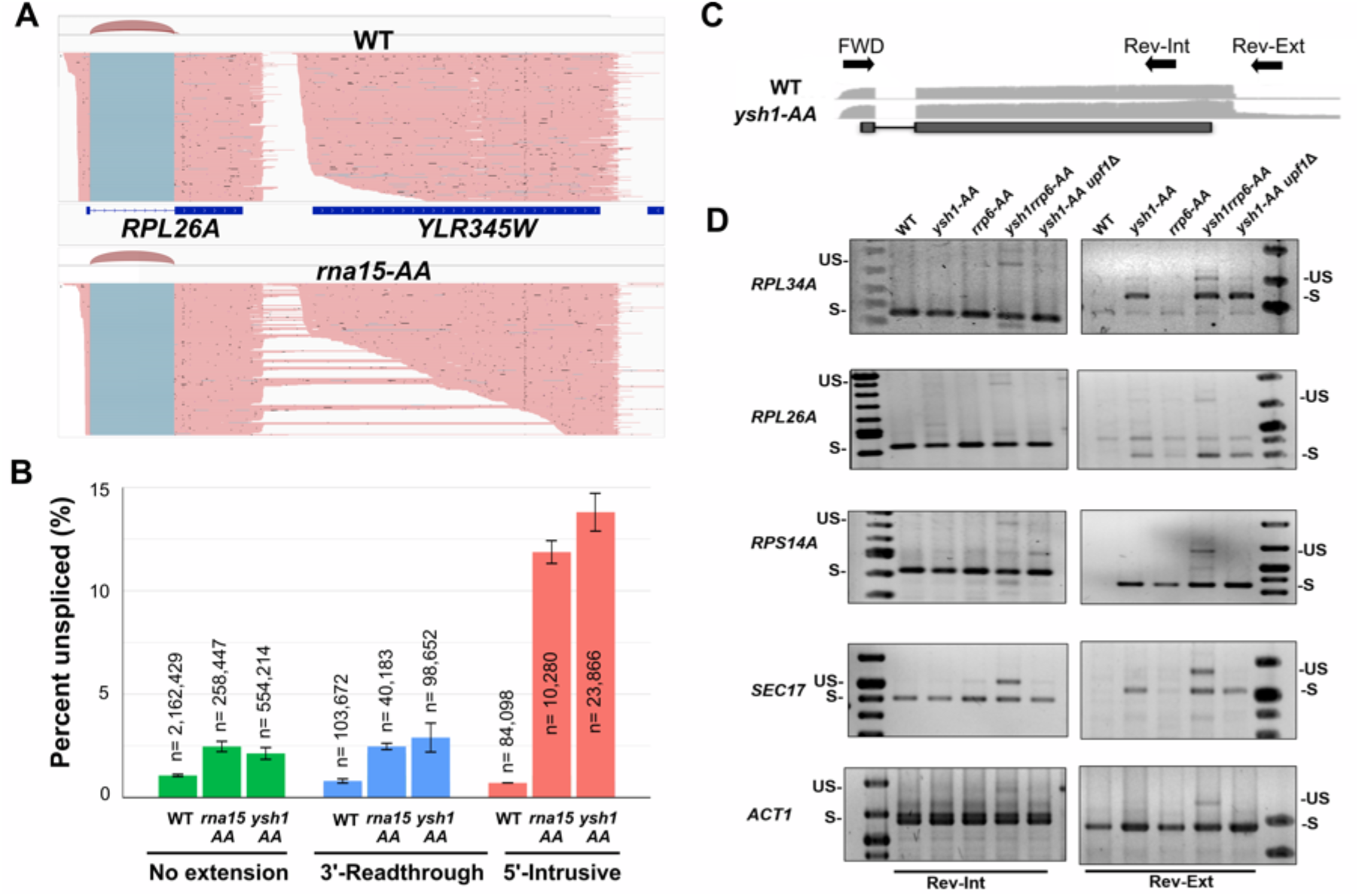
Impact of CPA inactivation on splicing efficiency. **A**. Selection of ONT reads from WT or Rna15AA in the *RPL26A-YLR345W* region showing the accumulation of 3′ extended reads of *RPL26A* in the *rna15-*AA strain. Reads are colored in salmon; splicing event connecting reads are colored in blue. Sashimi plots showing the splicing of *RPL26A* are also included. **B**. Bar graph showing the percentage of unspliced reads genome-wide for RNAs with no extensions, RNAs with 3′-readthrough, or 5′-intrusive in WT, *ysh1*-AA, or *rna15*-AA strains. Reads that contain a 5’ UTR that is 100 nt longer than what the annotated transcription start site (TSS) reported (36) were classified as 5′-intrusive. Reads that terminate more than 150 bp downstream of the annotated PAS were classified as 3′-extended/readthrough. Number in the bars indicate the number of sequenced reads included in each group. The p-values obtained for the comparisons are: *rna15*-AA vs WT: No Ext: 0.005848; *rna15*-AA vs WT: 3’ readthrough: 0.003757; *ysh1*-AA vs WT: No Ext: 0.007634; *ysh1*-AA vs WT: 3’ readthrough: 0.01342. **C**. RT-PCR scheme used to analyze the splicing status of normal and 3′-readthrough mRNAs. Shown is the Oxford Nanopore sequencing read coverage obtained for the WT and *ysh1*-AA samples for one ICG gene and the approximate location of the primers used for the RT-PCR analysis. The Rev-Ext primer only anneals to 3′-extended forms detected upon Ysh1 or Rna15 inactivation. Similar primer locations are used for all the genes analyzed. **D**. RT-PCR analysis of splicing of normal and 3′-extended mRNAs. Each panel shows RT-PCR products obtained from each indicated strains for the genes indicated. The Rrp6 protein was anchored-away (AA) for nuclear exosome inactivation (*rrp6*-AA). The *UPF1* gene was deleted (*upf1*<) for NMD inactivation. S=spliced; US = unspliced. Lanes on the right or left side of the gels with no label on top correspond to size markers.

The lack of effect of defect in 3′-end processing on splicing efficiency described above was surprising considering that nascent RNAs with 3′-extensions caused by Nab2 inactivation exhibited a high rate of intron retention (17). In contrast to this study which analyzed nascent transcripts using cDNA ONT sequencing, the ONT direct RNA sequencing platform used in our study relies on the presence of poly(A) tails for sequencing. For this reason, mRNAs that are 3′-extended but unpolyadenylated were undetectable in the sequencing experiments described above. Therefore, we could not rule out that unpolyadenylated, 3′-extended readthrough transcripts might be predominantly unspliced when the CPA machinery is inactivated. In addition, we could not rule out that RNA degradation pathways such as the nuclear exosome, or nonsense-mediated decay (NMD) might degrade unspliced transcripts, limiting their detection by Nanopore sequencing. To investigate the splicing status of unpolyadenylated, 3′-extended readthrough transcripts, we analyzed selected transcripts that did not show prevalent splicing defects in 3′-extended Nanopore reads by RT-PCR using total RNAs, which include unpolyadenylated RNAs. Two different reverse primers were used for each gene: an internal reverse primer (Rev-Int) that detects normal mRNAs, as well as 3′-extended ones; or a Rev-Ext primer that anneals only to 3′-extended RNAs (Fig2C). Concomitant inactivation of the nuclear exosome component Rp6p by anchor away, or deletion of the gene encoding the NMD factor Upf1p was used to assess if unprocessed transcripts are subject to nuclear or cytoplasmic surveillance. As show in Fig2D, RT-PCR analysis using the Rev-Ext primers showed that 3′-extended transcripts are readily detected in the *ysh1*-AA strain but absent in WT (or present at much lower level for *ACT1*). Importantly, this analysis did not reveal any major accumulation of unspliced 3′-extended transcripts in total RNAs extracted from the *ysh1*-AA strain (Fig2D), showing that defective 3′-formation does not promote major splicing defects even when analyzing bulk total RNAs. For all genes analyzed, a minor accumulation of unspliced 3′-extended transcripts could be detected upon inactivation of both Ysh1 and Rrp6. This result shows that a fraction of 3′-extended RNAs fail to splice and may correspond to primary transcript precursors which are efficiently degraded by the nuclear exosome, in agreement with prior studies(24)(25). These data are also consistent with the observation that unspliced and 3′-extended RNAs were detected upon sequencing nascent RNAs in the WT or *nab2*-AA strain(17), as sequencing of nascent RNAs is less sensitive to degradation pathways. By contrast, inactivation of NMD did not result in an increase in unspliced RNAs detection, which shows that these transcripts are likely retained in the nucleus, perhaps due their lack of poly(A) tails. A similar minor accumulation of unspliced transcripts was observed with the Rev-Int primer in the *ysh1*-AA/*rrp6*-AA strain. We note that concurrent inactivation of CPA factors and of Rrp6p has been shown to induce relocation of mRNAs to discrete nuclear compartments (26), which may complicate the interpretation of these data. Nevertheless, we conclude that unprocessed (unspliced and 3′-extended) RNAs do not accumulate to detectable levels unless the nuclear exosome is inactivated, and that the level of accumulation of these unprocessed transcripts varies depending on the genes analyzed (Fig2D).

### Transcription termination defects result in length-dependent splicing defects for downstream intron-containing genes

Analysis of the genes showing significant splicing defects in the *ysh1*-AA or *rna15*-AA samples revealed that unspliced reads were found predominantly in RNAs with 5′-extensions resulting from readthrough transcription from upstream genes (Fig2B). These 5′-extensions (defined as longer than 100nt than the annotated 5′-UTR, and referred to as resulting from ‘intrusive transcription’, as previously defined in (17)) are due to deficient termination from upstream transcription units, resulting in transcripts containing the upstream mRNAs, the intergenic region and the downstream ICG mRNA. One example is shown on Fig3A where deficient termination of *MRPL24* transcription generates intrusive transcripts into the downstream ICG *RPL36A*, leading to intron retention (Fig3A). This effect was detected genome-wide, as shown by a large increase in the fraction of unspliced reads for intrusive transcripts found in the *ysh1*-AA or *rna15*-AA strain relative to unextended transcripts found in any of the strains (Fig2B). This result is consistent with the data obtained on nascent transcripts upon Nab2 nuclear depletion (17), which resulted in the production of intrusive transcripts with a high rate of intron retention. Significantly, our data show that this effect can also be detected genome-wide on bulk RNAs when transcription termination is inactivated through inactivation of *bona fide* CPA factors.

The results described above suggested that an increased distance from the 5′-end of the transcript to the intron is detrimental to splicing. To provide support for this hypothesis, we compared the length of 5′ extensions found in intrusive transcripts showing either a significant splicing defect, or no splicing defects. In the *ysh1*-AA strain, the median 5′ extension size for ICGs which showed a significant splicing defect was 1320 nt, while the median 5′ extension size for ICGs which did not show a significant splicing defect was only 768 nt. In the *rna15*-AA strain, the corresponding numbers were very similar, at 1426 nt and 833 nt, respectively. Binning reads obtained from the *ysh1*-AA or *rna15*-AA strains according to 5′ extension length showed a threshold pattern, where transcripts with 5′ extensions smaller than 800 nt exhibited low levels of intron retention, while transcripts with longer 5′ extensions had significantly stronger splicing defects (Fig3B). We note that some direct RNA reads terminate short of the actual 5’ ends, which may have impacted the quantitative analysis of the impact of 5’-extension lengths on splicing efficiency. In particular, the splicing efficiency of RNAs with short 5’-extensions may have been underestimated, as a fraction of the reads assigned to the shorter extension groups may be the product of abortive sequencing of longer reads. Nevertheless, these data demonstrate that long 5′ extensions that are caused by readthrough transcription upon CPA inactivation decrease splicing efficiency of downstream ICGs.

The previous data showed that loss of termination due to nuclear depletion of CPA factors led to inefficiently spliced transcripts with 5′-extensions. To determine if this effect was solely due to the length of the 5′-exons/extensions or to indirect effects from CPA factors inactivation, we analyzed ONT data from WT cells to search for ICGs exhibiting reads with heterogenous 5′-ends and analyzed their splicing efficiency as a function of 5′-exon length. The *GIM4* ICG is located just downstream of the *YEA4* gene, and we detected a high number of transcripts originating from *YEA4* and reading through *GIM4* in WT sequencing samples (Fig3C). Strikingly, intrusive transcripts from *YEA4* showed a much lower splicing efficiency than the mRNAs originating from the bona fide *GIM4* promoter (Fig3C) as the level of unspliced reads for intrusive transcripts reached close to 50% (Fig3C). Similarly, the *YDL012C* gene showed transcripts heterogeneity in 5′-exon length, possibly due to usage of alternative transcription start sites (FigS5). Similarly to *GIM4, YDL012C* mRNAs with naturally long 5′-exons (>100nt longer than the location of the annotated transcription start-site) were less efficiently spliced than those with short 5′-exons (containing a 5’-UTR less than 100nt longer than the annotated one), with an increase of ∼5-fold in the amount of unspliced reads (FigS5). We note that the numbers shown on this figure might be an underrepresentation of the actual splicing defects for intrusive transcripts, as unspliced mRNAs can be targeted by various RNA surveillance systems (23) and as mentioned earlier, because of the issue of abortive sequencing for RNA with long 5’ -extensions.

To investigate whether long 5′-exons interfere with splicing efficiency in an orthogonal context, we designed two synthetic genes which express the *ACT1* gene with its intron and which differ by their 5′-exon length (Fig3D). One version includes an additional short sequence derived from the *PDC1* gene added to the *ACT1* 5′-exon, while the other includes the entire *PDC1* open-reading frame (Fig3D). RT-PCR analysis of WT cells expressing these constructs showed that the short 5′-exon version of *PDC1-ACT1* was spliced efficiently, while increasing 5′-exon length resulted in the detection of unspliced RNAs (Fig3D). Overall, the data obtained from quantitative analysis of long-read sequencing from wild-type and *ysh1*-AA or *rna15*-AA strains, as well as from the synthetic genes described above demonstrate that long 5′-exons interfere with splicing efficiency. The most direct explanation for this effect is that long 5′-exon length increase the distance from the 5′-cap to the intronic sequences, decreasing the activation of splicing by the cap-binding complex (5).

### Deficient transcription termination reveals dormant splice sites and promotes new splicing events

We next hypothesized that RNA extensions caused by loss of termination upon CPA inactivation would expose sequences matching splicing signals, which are otherwise not used in the context of normal termination but would result in novel splicing events. Inspection of the *ysh1*-AA and *rna15*-AA strain datasets identified several new splicing events. We detected frequent usage of a novel intron in *RPS16B*, where a cryptic 5′SS found in the second exon of *RPS16B* was spliced to a new 3′ SS in the *RPS16B* 3′UTR. This splicing event was detected in the *ysh1*-AA or *rna15*-AA strains, but not in WT (Fig4A; FigS6). The normal 5′SS of *RPS16B* was also found to be spliced to that new 3′SS (blue arrow, Fig4A) in the *ysh1*-AA or *rna15*-AA strains. It is possible that these cryptic splice sites are naturally used in a wild-type context. However, this would result in a very short transcript considering the length of the exons, which would likely be undetectable by Nanopore sequencing given the size limit for detection.

**Figure 3.**
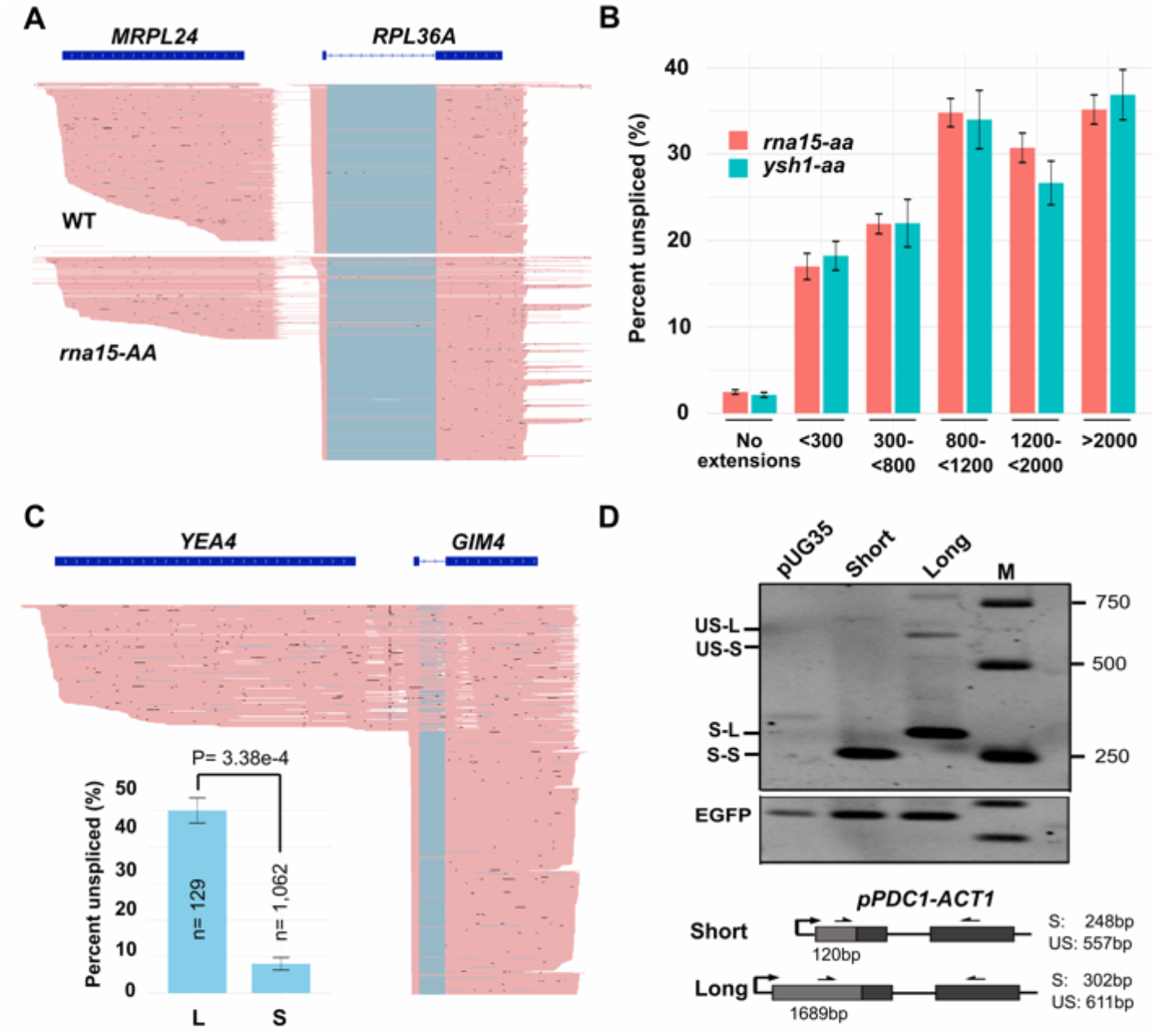
Intrusive transcription and long 5′ extensions inhibit splicing of downstream intron-containing genes. **A**. Selection of ONT reads from WT or *rna15*-AA in the *MRPL24*-*RPL36A* region showing the accumulation of unspliced 5′-extended reads of *RPL36A* in the *rna15*-AA strain. **B**. Bargraph showing the percentage of unspliced reads genome-wide as a function of 5′ extension size in the *ysh1*-AA or *rna15*-AA strains. Number in the bars indicate the number of sequenced reads included in each group. The p-values obtained for the comparisons are: *ysh1*-AA reads: <300 vs 300-800: 0.01846396591; <300 vs 800-1200: 0.01486; <300 vs 1200-2000: 0.03149; <300 vs >2000: 0.009079. *rna15*-AA reads: <300 vs 300-800: 0.007650383166 ; <300 vs 800-1200: 0.001137586334; <300 vs 1200-2000: 0.003206124344; <300 vs >2000: 0.004785227785. **C**. Selected Nanopore reads from WT control cells in the *YEA4-GIM4* region and quantification of unspliced RNA levels for *GIM4* mRNAs with short (S) 5′UTR, or readthrough mRNAs from the *YEA4* gene (L). Reads at the top of the picture are intrusive transcripts from *YEA4* gene into *GIM4* which show a high fraction of unspliced reads; Numbers on the y-axis indicate the percentage of unspliced reads. **D**. Synthetic *PDC1-ACT1* gene versions and RT-PCR analysis of splicing. Shown are RT-PCR products obtained from RNAs extracted from WT strains transformed with the empty vector (pUG35) or pUG35 plasmids expressing the short or long 5′ exon version of the *PDC1*-*ACT1* gene fusion depicted, using the forward (Fwd) and reverse (Rev) primers indicated on the figure. RT-PCRs for the empty vector sample were performed using the short *PDC1-ACT1* primer pair. The eGFP transcript expressed from the pUG35 vector was used as a loading control. M indicate the lane with the size marker.

**Figure 4.**
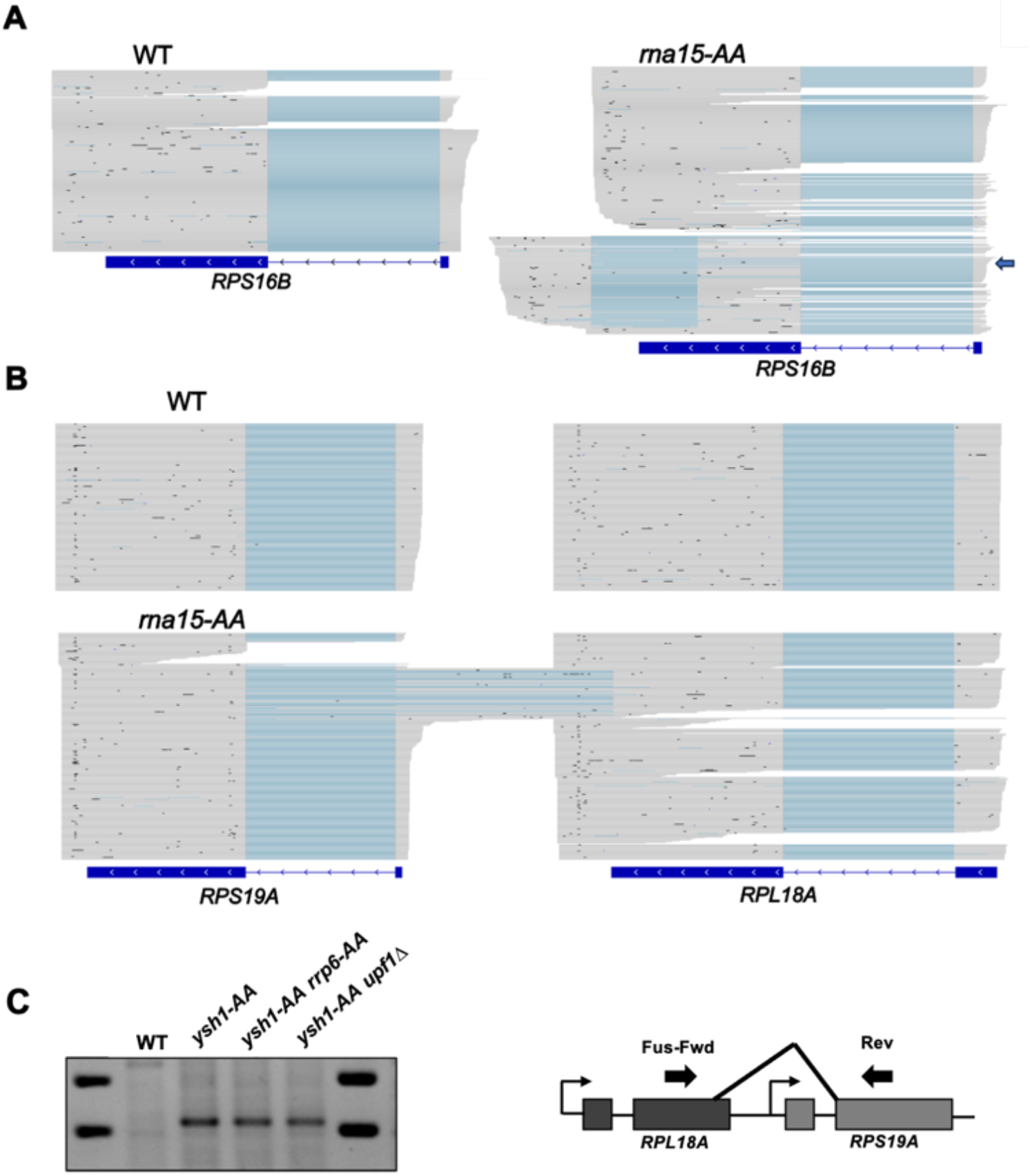
Novel Splice sites are activated by deficient transcription termination. **A**. Selection of Nanopore sequencing reads for *RPS16B* in the WT control or *rna15-AA* strain. For *rna15-AA*, two groups of reads are presented. The reads group at the top shows splicing patterns similar to WT. The read group at the bottom shows usage of the cryptic 5′ and 3′ SS in exon2 of *RPS16B*. The blue arrow indicates reads for which the normal 5′ SS of *RPS16B* is spliced to the cryptic 3′ SS. **B**. Selection of Nanopore sequencing reads for the *RPL18A* and *RPS19A* genes region for WT and *rna15-AA* showing the intergenic *RPL18A-RPS19A* splicing using the new 5′SS found at the end of *RPL18A*. **C**. RT-PCR detection of intergenic *RPL18A*-*RPS19A* splicing. RNAs extracted from WT control or *ysh1*-AA,*ysh1*-AA/*rrp6*-AA, and *ysh1*AA-*upf1*< were analyzed by RT-PCR using the forward (Fwd) and reverse (Rev) primers indicated on the figure.

In another case, we detected frequent usage upon Ysh1 or Rna15 inactivation of a cryptic 5′SS at the end of the second exon of *RPL18A* spliced to the 3′SS of the downstream gene *RPS19A* (Fig4B). The GUAAGU sequence of this cryptic 5′SS overlaps with the UAA stop codon of *RPL18A* (FigS7). This splicing event resulted in the production of chimeric *RPL18A-RPS19A* transcripts, which corresponds to about 10% of the splicing events for extended species of *RPL18A* in *rna15*-AA or *ysh1*-AA. (Fig4B). We validated the production of this spliced product by RT-PCR in the *ysh1*-AA strain (Fig4C). Interestingly, this spliced isoform did not appear to be targeted by the nuclear exosome (*ysh1*AA *rrp*6AA, Fig4C), nor by nonsense-mediated decay (NMD) (*ysh1*-AA-*upf1τ*<, Fig4C). This chimeric mRNA would encode the Rpl18 protein in its entirety, fused to the protein sequence encoded by the second exon of *RPS19A*. In the case of the duplicated *RPL18B* gene paralogue, the corresponding sequence is GUAAUC and we did not detect splicing between *RPL18B* and *RPS19B* at that site. The mismatches at positions 5 and 6 compared to canonical 5′SS sequences explains why such intergenic splicing events were not detected for the paralogue genes.

### Deficient transcription termination promotes long-distance intergenic splicing events, including between adjacent splicing-dormant genes

The previous example showed that loss of termination could promote the production of chimeric *RPL18A*-*RPS19A* transcripts. In *S*.*cerevisiae*, several intron containing genes (ICG) are located in tandem on the same strand (Table S3). Deficient termination due to inactivation of Ysh1 or Rna15 may therefore result in the production of polycistronic transcripts containing multiple introns, providing the opportunity for splicing between adjacent genes. We searched for such intergenic splicing events by identifying reads with splicing junctions between sequences belonging to different genomic features. After curating these junctions for annotated overlapping transcription units and read alignment artefacts, we identified several cases of intergenic splicing triggered by loss of termination in the *ysh1*-AA and/or *rna15*-AA strains. The most frequent event detected was the splicing of the 5′SS of *RPL22B* to the 3′SS of the downstream gene *MOB2* (Fig5A). It was shown previously that an alternative intronic 5′ SS is present in *RPL22B* (27). This alternative 5′SS was also found to be spliced to the downstream 3′ SS of *MOB2*. Reads showing splicing of exon1 of one ICG to the exon2 of a downstream ICG were also detected for several other gene pairs: *RPL18B-RPS19B, RPS9A-RPL21B* and *RPS24A-YOS1* (Fig5B). These cases of intergenic splicing were detected upon inactivation of either Rna15 or Ysh1, showing that deficient transcription termination can promote intergenic splicing regardless of which factor is inactivated. In addition to the events described above where natural splice sites of adjacent genes are used, we also discovered cases where dormant splice sites are revealed. In the case of *IDP1*, a cryptic 5′ SS is present at the beginning of the ORF, which is spliced to the 3′ SS of the downstream *UBC9* gene (Fig5B; FigS8). Strikingly, in the case of the *LCB2-AIM7* or *DUT1-SRB6* gene pairs, none of these genes are naturally spliced. However, Ysh1 or Rna15 inactivation generates readthrough species which results in the intergenic splicing between a 5′SS present in the 5′UTR or ORF of the upstream gene (*LCB2* or *DUT1*) and a 3′ SS in the ORF of the downstream gene (*AIM7* or *SRB6*, respectively; Fig5B; FigS9). We used RT-PCR to validate two of these splicing events (Fig5C), and test whether these intergenic spliced species are subject to nuclear surveillance or degradation by NMD. In agreement with the Nanopore sequencing data, intergenic splicing between *RPL22B* and *MOB2* or between *DUT1* and *SRB6* was detected only upon inactivation of Ysh1, but not in the WT control (Fig5C). There was no major effect of inactivating Rrp6 or the NMD factor Upf1 on the accumulation of these intergenic spliced products, suggesting that these species are not subject to these RNA degradation systems.

**Figure 5.**
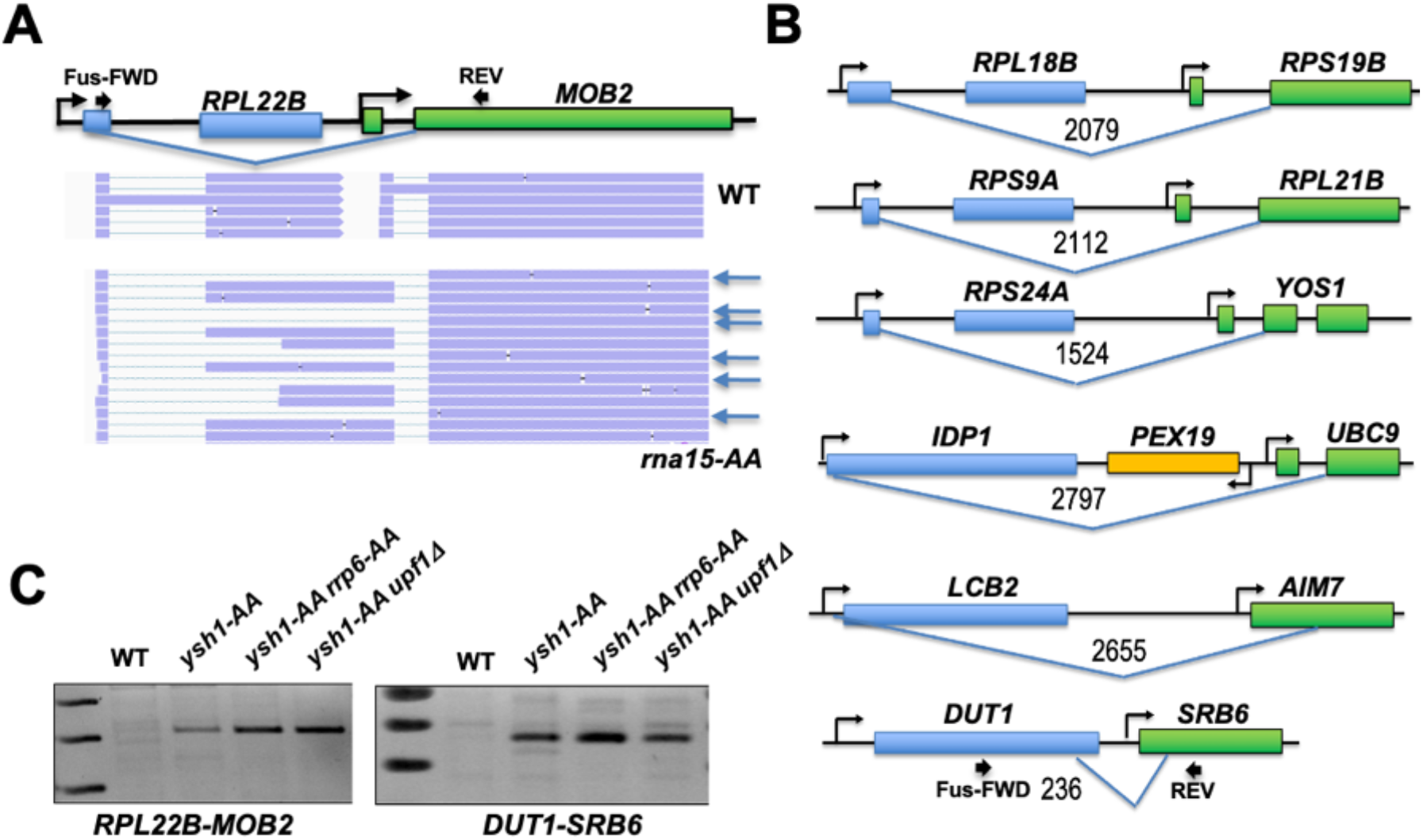
Deficient Termination promotes Intergenic splicing. **A**. *RPL22B*-*MOB2* genomic region and examples of Nanopore sequencing reads obtained from WT or *rna15-AA* strains. Reads highlighted with blue arrows indicate intergenic *RPL22B*-*MOB2* splicing events. **B**. Other examples of intergenic splicing events detected in the *ysh1*-AA and/or *rna15-AA* strains. Numbers indicate the distance between the 5′ and 3′ SS used. **C**. RT-PCR analysis of intergenic splicing in WT, *ysh1*-AA and derivative strains. The approximate locations of the forward (Fwd) and Reverse (Rev) primers used is indicated on panel (A) for *RPL22B*-*MOB2* or (B) for *DUT1*-*SRB6*.

## Discussion

In this work we have studied the effect of inactivating splicing or 3′-end formation on splicing and 3′-end formation using long-read ONT sequencing. The use of this platform allowed us to directly investigate the splicing efficiency and 3′-end of transcripts genome-wide. Strikingly, we did not find evidence for global reciprocal coupling between splicing and usage of the adjacent downstream 3′-end processing signals. Splicing inactivation did not perturb 3′-end processing (Fig1), while inactivation of CPA had a limited impact on splicing, which was mainly due to intrusive transcription, as previously shown upon nuclear depletion of Nab2(17). One of the limitations of our work is that ONT sequencing relies on polyadenylated RNAs, and RNAs showing 3′-extensions in our sequencing data are still polyadenylated albeit at more distant sites. However RT-PCR analysis from total RNAs (which include both polyadenylated and unpolyadenylated RNAs) of selected transcripts showed that 3′-extended transcripts detected upon Ysh1 inactivation are mainly spliced and that a minor population of unspliced and 3′-extended transcripts can be detected if the nuclear exosome component Rrp6 is inactivated concomitantly with Ysh1 (Fig2D). The involvement of the RNA exosome in degrading unprocessed RNAs is in agreement with previous studies showing that readthrough transcripts generated upon genetic inactivation of some CPA factors can be degraded by the exosome (24). While the detection of unspliced and 3′-extended transcripts upon inactivation of a CPA factor and of the nuclear exosome may suggest some level of coupling of splicing to 3′-end processing on a transcript-specific basis, we note that the majority of 3′-extended transcripts are still efficiently spliced (Fig2D). Moreover, simultaneous inactivation of CPA factors and of Rrp6p has been shown to induce relocation of mRNAs to discrete nuclear compartments (26), which may complicate the interpretation of data involving these double mutants. Based on our results, we conclude that the splicing process has little effect on 3′-end processing, and that there is no global coupling of splicing to recognition of the immediate polyadenylation sites for most *S*.*cerevisiae* ICGs, although coupling might exist for a few transcripts such as *RPL28* (FigS4). One reason for the loss of a general coupling mechanism is that unlike in mammalian genomes, most transcription units in *S*.*cerevisiae* do not contain intronic sequences. Therefore the evolutionary pressure to maintain functional coupling between these two processes may have been reduced if the signals that dictate the efficiency of each process are strong enough.

One of the major conclusions from our work is that defective transcription termination results in intrusive transcripts with decreased splicing efficiency, as initially reported through sequencing of nascent RNAs upon nuclear depletion of the nuclear poly(A) binding protein Nab2(17). While Nab2 is not a bona fide component of the CPA machinery, it is possible that the global impact of nuclear depletion of Nab2 on nuclear RNA decay (28) and/or on mRNA export (29) indirectly impairs transcription termination and/or CPA. We also note that the effect of Nab2 inactivation on splicing was detected primarily for ICGs with suboptimal splicing signals(17), and that Nab2 was found to crosslink to intronic signals(30), which suggests that the results obtained upon Nab2 inactivation may be due to a combination of effect on splicing and transcription termination. By contrast, we did not observe a bias in suboptimal splicing signals for RNAs which show defective splicing upon Ysh1 or Rna15 inactivation (Table S2), suggesting that the splicing defects we detected are solely due to termination defects. Interestingly, the deleterious impact of long 5′extensions on splicing can also be detected in normal cells for transcripts exhibiting naturally heterogeneous 5′-ends such as *GIM4* (Fig3C) or *YDL012C*, or for synthetic constructs with a long 5′-exon (Fig3D). These results show that the splicing defects induced by long extensions are not intrinsically due to the absence of CPA factors but suggest that an increase distance of the splicing signals to the 5′-cap structure is responsible for the impairment of splicing, as it would decrease the positive effect of the cap-binding complex on splicing. Alternatively, we cannot rule out that long 5′-extensions might promote secondary structures that impair the recognition of downstream splicing signals. However, given the general widespread effect of 5′ extension length on splicing, we believe this hypothesis to be unlikely. Regardless of the specific mechanism by which long 5′-exons interfere with splicing, these observations suggest that the frequent occurrence of short 5′-exons in *S*.*cerevisiae* ICGs is likely to result from the evolutionary pressure to prevent the potential deleterious effects of long 5′-exons on splicing. Overall our findings, combined with those obtained with Nab2(17) underscore the importance of proper termination to limit readthrough transcription onto downstream genes, which is particularly critical in a compact genome such as *S*.*cerevisiae*.

One of the most striking effects of inactivating CPA is the production of polycistronic transcripts with multiple splicing signals, resulting in the detection of novel splicing events (Fig4, 5). These results provide further evidence that inhibition of CPA does not impair splicing, as the extended species that are produced upon CPA inactivation increase the repertoire of splicing events detected. Intergenic splicing events can be highly abundant in the case of *RPL22B-MOB2* or rare, as they are sometimes detected with only a few reads in the *ysh1*-AA or *rna15*-AA samples. Intergenic splicing events might be underrepresented in these ONT RNA sequencing datasets for two major reasons. First some of these species might be targeted by RNA surveillance pathways, even if we did not detect a major effect upon inactivating the nuclear exosome or NMD (Fig4,5). For instance, exon skipped products (which functionally correspond to the intergenic spliced products we describe here) have been shown to be targeted by the exosome and Rat1 in *S*.*cerevisiae* (31). In addition, 5′-exons of *S*.*cerevisiae* ICGs can be very short, and the 3′-end bias observed in Nanopore sequencing data from yeast (32) may result in the underrepresentation of spliced transcript with short 5′-exons. We note that the distance involving some of these splicing events exceeds the size of the largest natural intron in *S*.*cerevisiae*, including several detected splicing events involving splice sites that separated than more than 2kb, including 2.6 and 2.8kb (Fig5B). These results indicate that the yeast splicing machinery can operate at longer distances than previously thought. On the other hand, readthrough between closely positioned transcription units can promote short distance splicing events such as the one detected between *DUT1* and *SRB6*. It is possible that some of these intergenic splicing events may occur in natural conditions and produce functional products if natural levels of transcriptional readthrough occurs, such as in stress conditions, as reported in mammalian cells(33). The compactness of the yeast genome and the close spatial positioning of transcription units make this more likely and could potentially provide a strategy to produce proteins resulting from splicing of adjacent genes.

## Materials and Methods Yeast Strains and Growth

All yeast growth was performed in YPD media (1% w/v yeast extract, 2% w/v peptone, and 2% w/v

dextrose). In all experiments cells were back diluted from an overnight solution to an O.D.600nm between 0.1 and 0.13 in 50mL per time-point and grown for approximately two doublings at 30°C, shaken at 200rpm. 50mL of cell culture would be harvested via centrifugation and flash frozen in liquid nitrogen for downstream use. Anchor away experiments were performed by replacing cell culture media of log-phase yeast with YPD+ 1µg/mL of rapamycin as described (18). Cells were then grown for an additional 2 hours in YPD+rapamycin before being harvested by centrifugation and flash frozen for downstream use. Strains used in this work can be found in Table S4.

### Plasmids design and synthesis

The two plasmids expressing the short and long versions of the PDC1-ACT1 gene fusions were synthesized by Twist. For both constructs, 549bp upstream of the natural *PDC1* 5′ UTR and the 5′ UTR of *PDC1* were used. The “short” version includes the first 120 coding nucleotides of PDC1 fused to the start codon of *ACT1* up to 33bp after the 3′ UTR of ACT1, while the “long” version, contains the full *PDC1* open-reading frame (excluding the stop codon) fused to the same region of *ACT1* (see Fig3D). Because of synthesis issues with homopolymeric sequences, one change was made in *ACT1* where a 14bp long poly(A) intronic sequence from positions 189-202 of *ACT1* was changed to ATATAATTAATTAA. The PDC1-short_ACT1 and PDC1-long_ACT1 sequences were inserted into the empty pUG35 vector 49bp downstream of the lac operator sequence, and 64bp upstream of the MET17 promoter.

### RNA Extraction and Direct RNA sequencing

Total RNAs were extracted in triplicate via the standard phenol-chloroform technique(20) from anchor-away yeast strains. RNAs for the wild-type, *ysh1*-AA, and *rna15*-AA samples were then further processed according to the protocol described in the manufacturer′s manual for ONT product (SQK-RNA004) for sequencing on the PromethION system (PRO-SEQ002) using RNA Flow Cells (FLO-PRO004RA). The only exception was the use of Maxima-H Minus as the RT enzyme as this was the recommended enzyme at the time of these experiments. RNAs for the snp1-AA samples were further processed per the manufacturer′s manual for ONT product (SQK-RNA004) for sequencing on the MinION system (MIN-101B) using RNA Flow Cells (FLO-MIN004RA) using the current Induro RT enzyme protocol. RNA Flow Cells were washed according to the manufacturer′s protocol for the Flow Cell Wash Kit (EXP-WSH00) in between sample runs.

### RT-PCR Analysis

RNAs for RT-PCR were generated from yeast cells as described above. RNA samples were then DNase I treated according to the manufacturer′s instructions (Thermo Fisher EN0521). For cDNA synthesis from total RNAs the standard Maxima H Minus RT protocol (Thermo Fisher EP0751) was followed with gene specific primers (Table S5) and 100 units of enzyme per reaction. 1uL of each RT reaction was then used as template for a 20uL HiFi-PCR reaction using the corresponding gene specific primers (Table S5). 5uL PCR products were then mixed with a 6x Bromophenol Blue and Xylene Cyanol loading dye and analyzed by agarose gel using 1.2-1.8% w/v agarose in 1xTAE (40mM Tris Base, 20mM acetic acid, 1mM EDTA, pH 8.2-8.4) depending on expected product size. Gels were then visualized by Ethidium Bromide staining on an Azure 300 Imaging System.

### Bioinformatics and Statistical Analysis

Reads were basecalled in real-time using the default algorithm for direct RNA sequencing on MinKNOW (version 6.2.6) with Dorado (version 7.6.7). Reads were then mapped to the Saccharomyces cerevisiae genome (S288C_reference_sequence_R64-5-1) acquired from SGD using Minimap 2 (Version 2.17-r941). The following parameters were used during alignment “minimap2 -ax splice -G 3000” which set a maximum intron length of 3000 bp. Resulting bam files were coordinate sorted and converted to bed files using SAMtools(34) and Bedtools (https://bedtools.readthedocs.io/en/latest/) for downstream analyses. Reads were visualized using IGV (Version 2.12.3).

Reads were annotated with overlapping genes using bedtools intersect (v2.30.0). For read counts, we calculated the number of reads mapping to each gene using featureCounts as part of the Rsubread package (version 2.12.3) allowing for multiple overlaps. DESeq2(35) was used to quantify changes in gene expression and for normalizing counts to library size.

3’UTR lengths were calculated by subtracting the 3’-end of reads from the end position of their assigned gene open-reading frame (annotated stop codon). The median 3’ UTR length was calculated for each gene, and the average of the median3’ UTR length was calculated for all 3 replicates. For the analysis of splicing efficiency, counts were generated separately for whole genes and introns specifically. For a read to be counted as unspliced it needed to overlap an annotated intron by at least 6 bases. Fraction of unspliced reads was generated by dividing intronic read counts by overall gene counts. For extension analysis, reads were classified into 4 groups. “3 prime extended” are reads which overlap the gene of interest but terminate more than 150 bp downstream of the annotated PAS. “5 prime intrusive” overlap the gene of interest, but begin more than 100 bp upstream of the annotated TSS(36). If a read met both of these conditions, it was assigned as an “intrusive transcript”. All other reads fall into the “no extension” category. Experimentally derived UTR annotations(36) were used for the TSS and PAS locations.

All statistical tests and p-values reported were based on a welch’s t-test. For the splicing volcano plots the p-value was adjusted further using the Benjamini-Hochberg method to control for False Discovery Rate.

## Data availability

Custom Nanopore processing code is available on GitHub: https://github.com/kevinh97/Yeast-Transcription-termination. The long-read RNA sequencing data reported in this paper are accessible in the NIH SRA database under the accession number PRJNA1229592.

## Acknowledgments

We thank Douglas Black and members of the GFC lab for discussion and Alina Tong for comments on the manuscript. KH was supported by a Dissertation Year Fellowship from the UCLA Division of Graduate Education. Supported by NIGMS grant GM130370 to GFC.

