## Supplemental Figures and Tables for "Transcription termination promotes splicing efficiency and fidelity in a compact genome"

**Supporting Information– Barr, He et al.**

**Figure S1. Reproducibility of Oxford Nanopore Long Read RNA Sequencing data.** Plots showing normalized gene counts for each replicate among the 4 different strains sequenced (WT, YSH1-AA, RNA15-AA, and SNP1-AA). Counts were generated using FeatureCounts as part of the Rsubread package, DESeq2 was used for normalizing counts to library size.

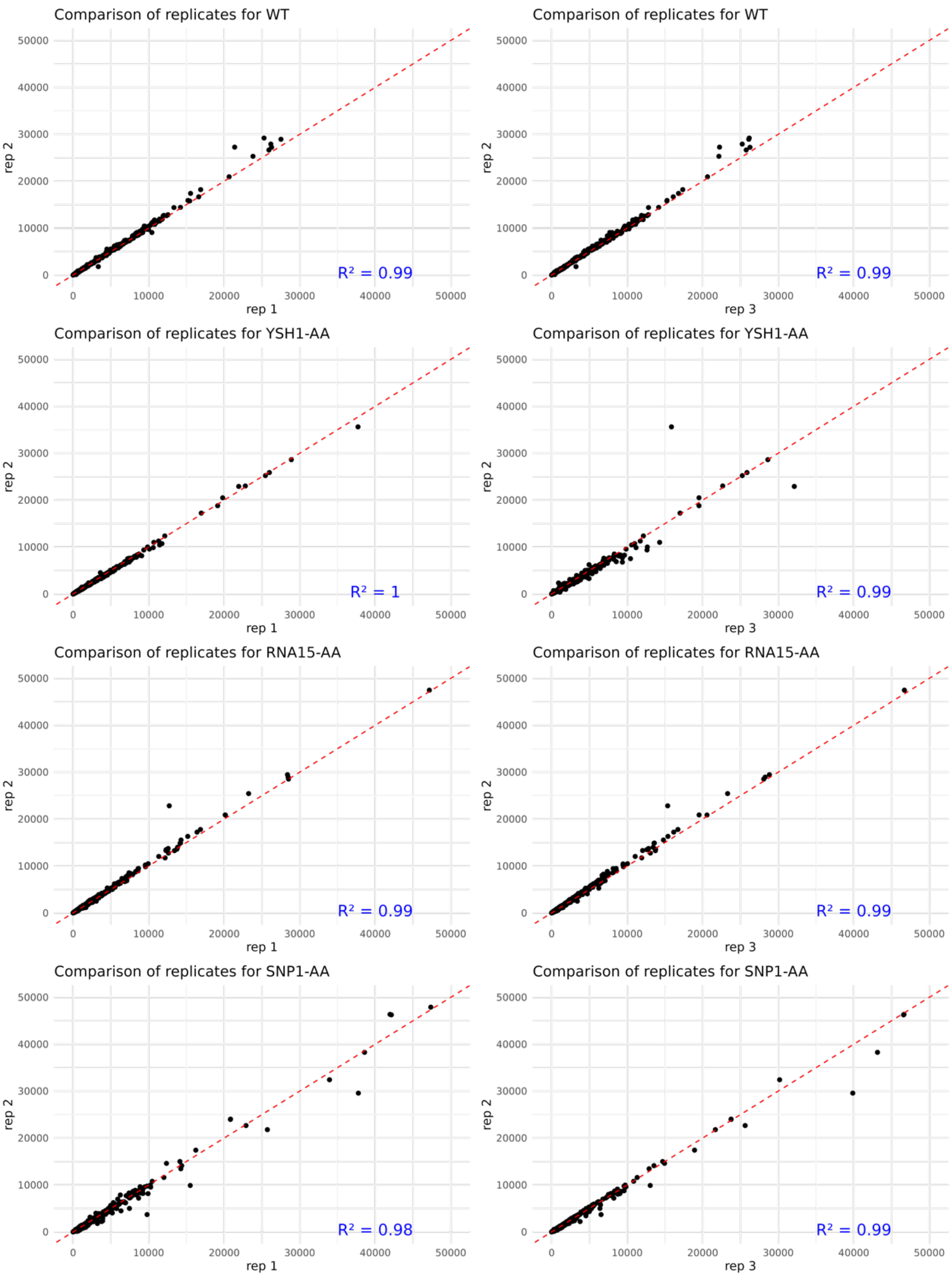

**Figure S2. Example of genes that show increased usage of distal poly(A) sites in the Snp1-anchor away.** Shown are 3′-end peaks detected for the control wild-type sample or the Snp1, Ysh1 or Rna15-anchor away strains in the 3′-UTR of *COX6* (A) or *RPB2* (B). The y-axis represents the 3′UTR region positions, with the origin corresponding to the end of the stop codon.

**A.** Distribution of COX6 poly(A) site usage in the indicated strains.

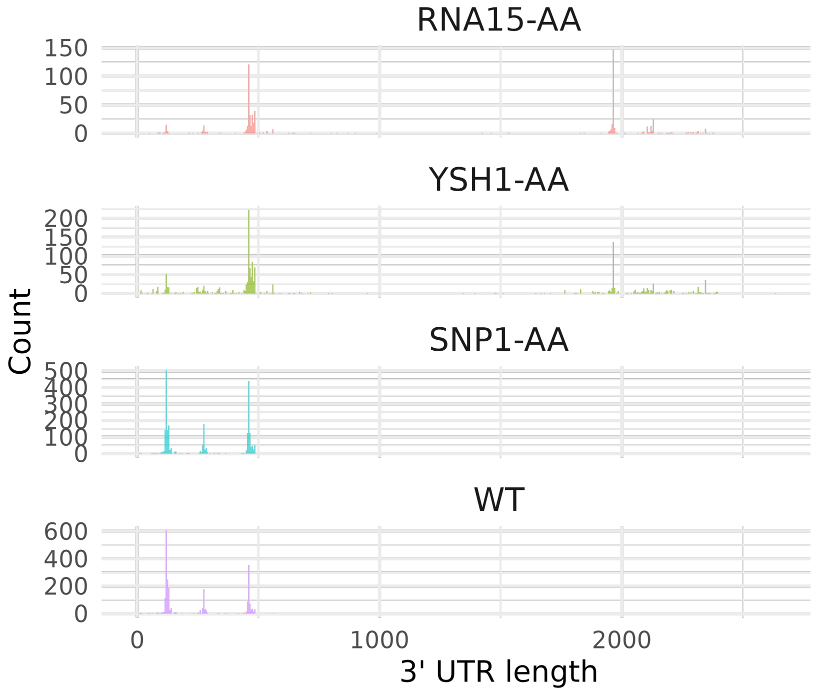


**B.** Distribution of RPB2 poly(A) site usage in the indicated strains.

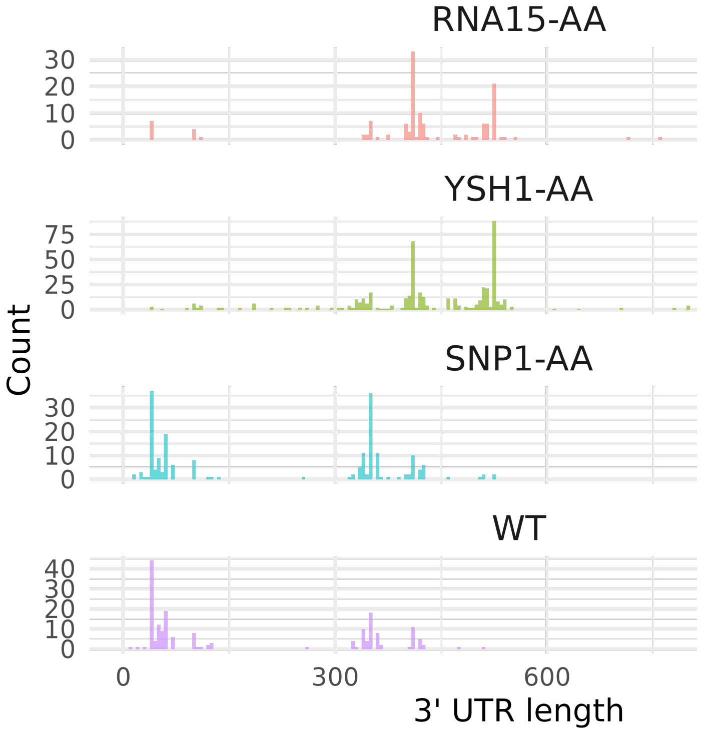

**Figure S3.** **S**equence logos in the 5' splice site or branchpoint regions of all intron-containing genes (ICG) or of mRNAs which show an increase of intron retention specifically in the Ysh1-AA or Rna15-AA strains.

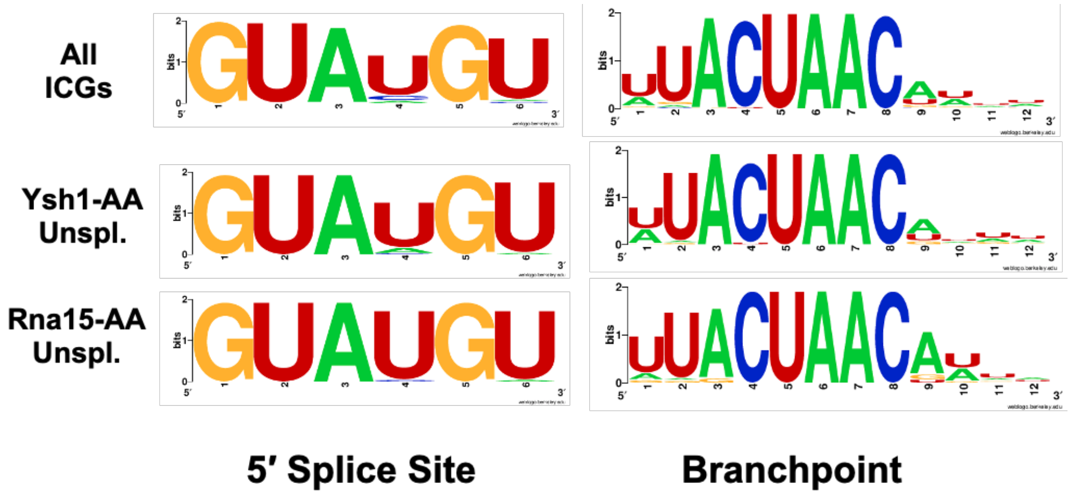


**Figure S4. Unspliced reads accumulation and intron-retention quantifications for *RPL28***
Shown is a selection of ONT reads from WT or Rna15-AA showing the accumulation of unspliced and 3′ extended reads of *RPL28* in the Rna15AA strain. The bargraph shows on the y-axis the percentage of unspliced reads in WT, Rna15-AA or Ysh1-AA for reads with no extensions or 3'extensions.

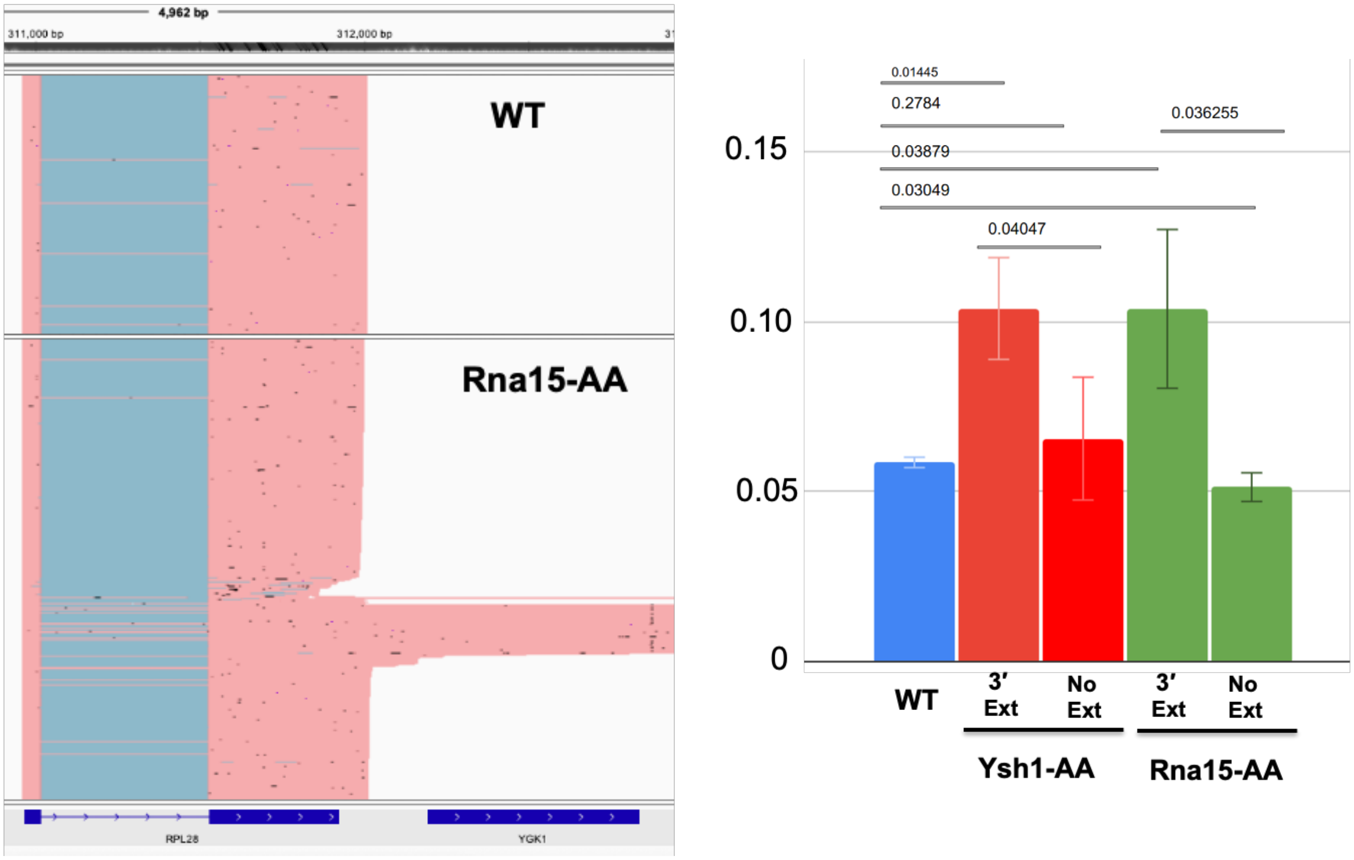

**Figure S5. Oxford Nanopore sequencing and intron-retention quantification for *YDL012C* in wild-type cells.**

Shown in a selection of Nanopore reads from WT control cells in the *YDL012C* region and the quantification of splicing defects for YDL012C mRNAs with short 5′-exons (S), or with 5′ exons at least 100nt longer than the annotated 5′ exon (L); Numbers on the y-axis indicate the percentage of unspliced reads. Numbers above each bar indicate the number of reads included in the analysis. p-value L vs S = 0.01879.

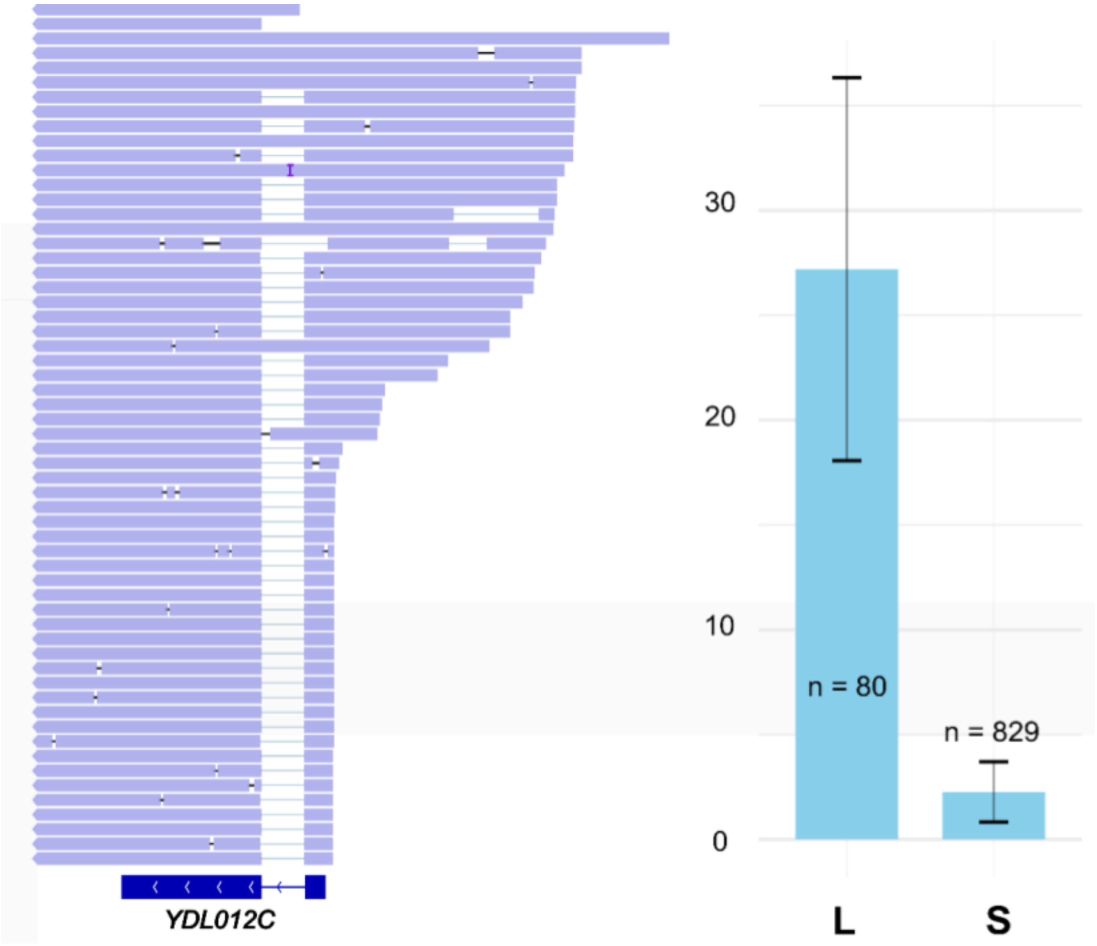

**Figure S6. A new intron is activated in *RPS16B* in the Ysh1-AA or Rna15AA strains.** Shown are sashimi plot of splicing events detected in the WT or Rna15AA strain as well as reads showing usage of the cryptic intron in the *RPS16B* 2nd exon. The 5′ SS located in the middle of exon2 and 3′ SS located in the 3′ UTR used are highlighted in red.

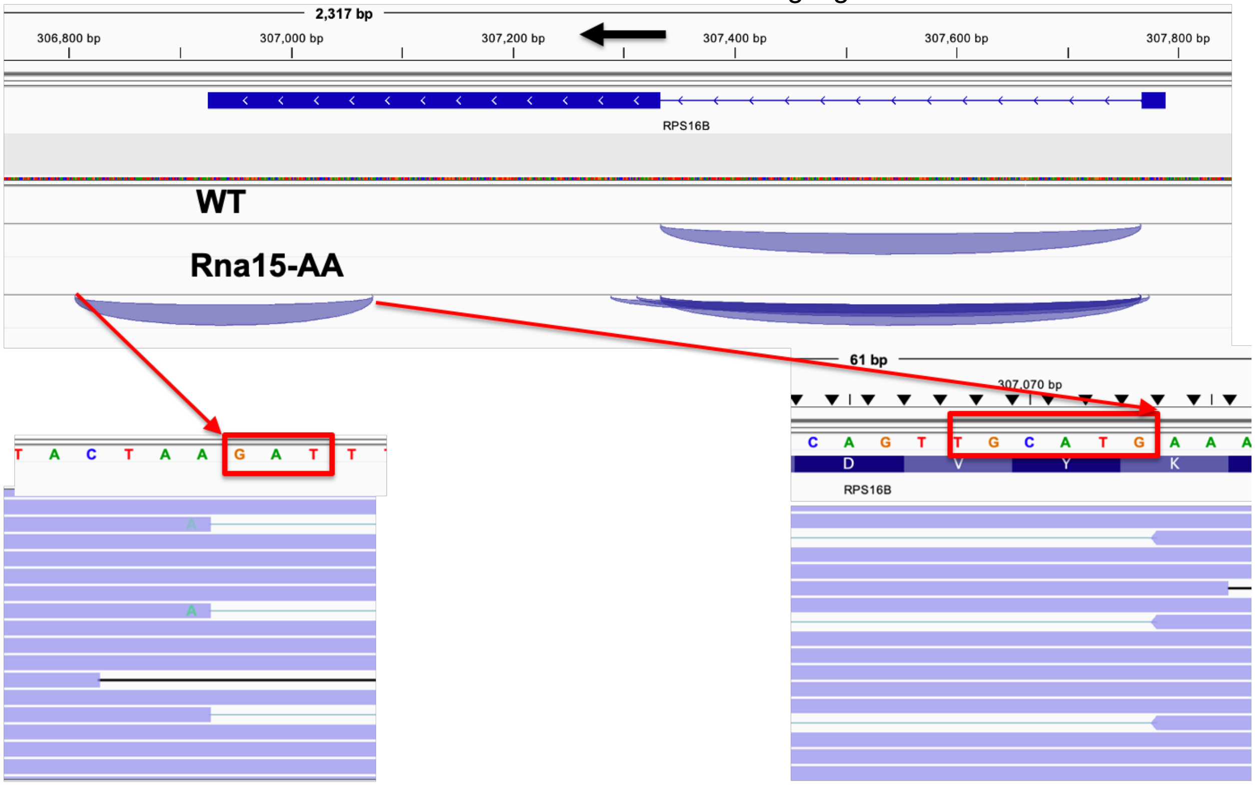


**Figure S7. A 5′SS used in the Ysh1-AA or Rna15-AA strains overlaps with the stop codon of *RPL18A*.**

Shown are sequencing reads from the Rna15-AA strain highlighting usage of the GUAAGU 5′ SS highlighted in red overlapping with the UAA stop codon of *RPL18A*. Reads polarity are from right to left since the gene is localized on the C (-) strand.

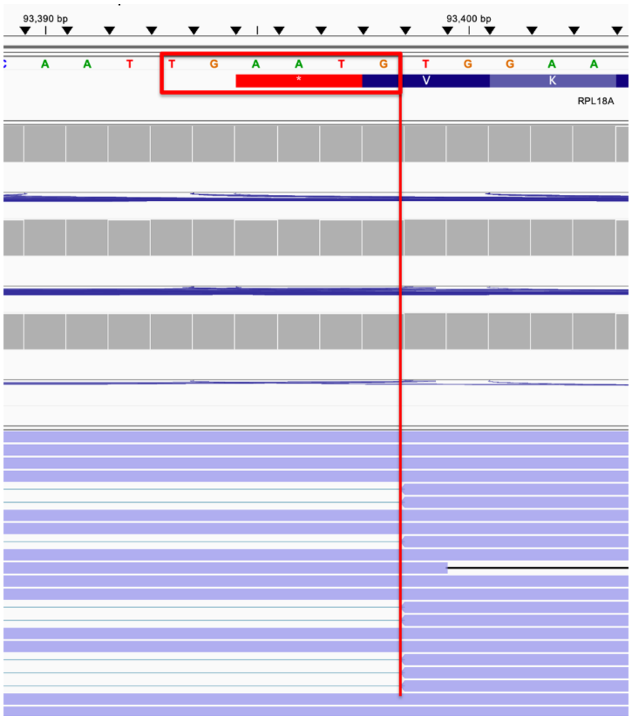

**Figure S8. *IDP1-UBC9* intergenic splicing events detected in the Rna15 or Ysh1AA strain.** Shown are reads obtained from WT or from the Rna15-AA strain showing the splicing between the 5′ SS at the beginning of *IDP1* and *UBC9*. The cryptic 5′ SS sequence at the beginning of the *IDP1* ORF is highlighted in red.

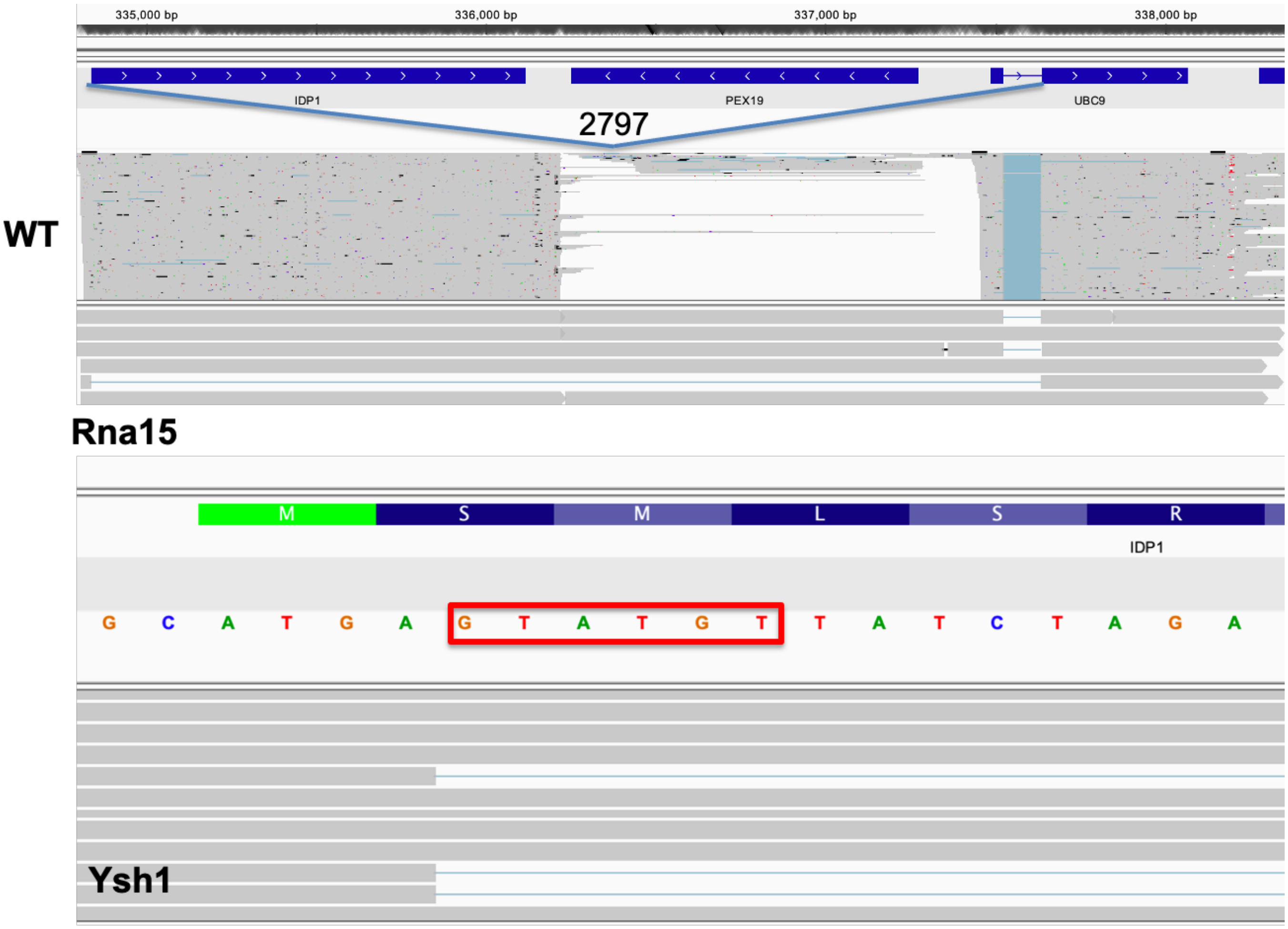

**Figure S9. *LCB2-AIM7* intergenic splicing events detected in the Rna15 or Ysh1AA strain.** Shown are reads produced by the LCB2-AIM7 intergenic splicing events detected in the Ysh1AA strain. Reads from WT samples are also included. The cryptic 5′ SS sequence in the *LCB2* 5′UTR and the putative branchpoint and 3′ SS used in AIM7 are is highlighted in red.

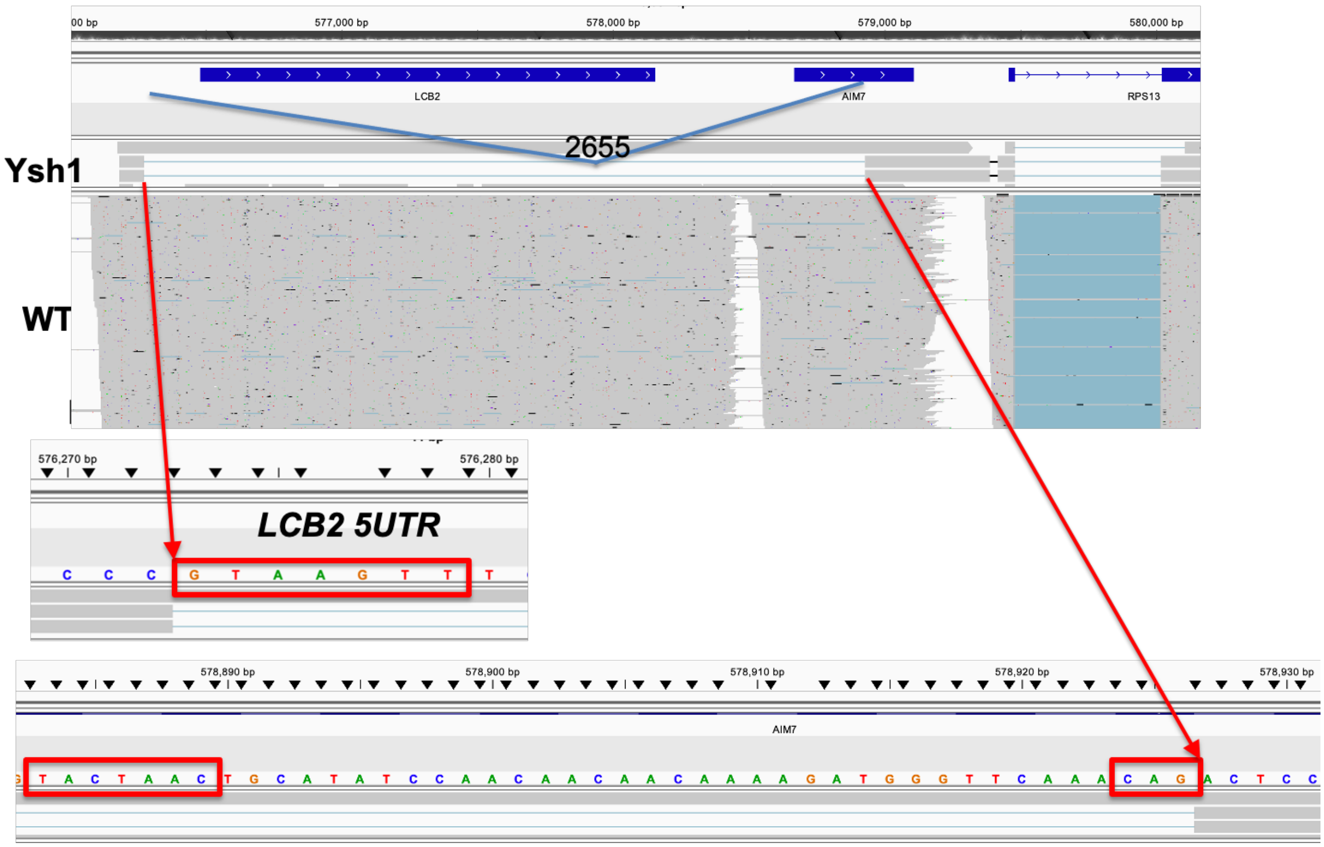

**Table S1**

Statistics of the number of reads obtained by Oxford Nanopore Sequencing for each strain and replicate samples. Also shown is the number of reads that aligned to the S.cerevisiae genome and the percentage of reads aligned.

| Strain | Total reads | Aligned reads | Percent Aligned |
|---|---|---|---|
| **WT rep1** | 4185196 | 3809153 | 91.014% |
| **WT rep2** | 4152656 | 3715956 | 89.483% |
| **WT rep3** | 2745690 | 2497176 | 90.948% |
| **RNA15 rep1** | 1052882 | 908599 | 86.296% |
| **RNA15 rep2** | 1835357 | 1593438 | 86.818% |
| **RNA15 rep3** | 915672 | 787530 | 86.005% |
| **YSH1 rep1** | 2780820 | 2358478 | 84.812% |
| **YSH1 rep2** | 4253350 | 3603440 | 84.720% |
| **YSH1 rep3** | 1787771 | 1556736 | 87.076% |
| **SNP1 rep1** | 3298600 | 2838622 | 86.055% |
| **SNP1 rep2** | 6436917 | 5471358 | 84.999% |
| **SNP1 rep3** | 2681055 | 2289439 | 85.393% |

### Table S2. List of genes that show extended median

| gene_name | WT_median_extensions | WT_gene_counts | Snp1_median_extensions | Snp1_gene_counts | deviation |
|---|---|---|---|---|---|
| ARN2 | 162 | 63 | 330 | 41 | 168 |
| CBK1 | 123 | 69 | 373 | 75 | 250 |
| COX6 | 129 | 2296 | 271 | 2300 | 142 |
| FEN2 | 723 | 107 | 258 | 124 | -465 |
| FZF1 | 201 | 75 | 644 | 111 | 443 |
| GTB1 | 167 | 66 | 293.5 | 78 | 126.5 |
| IRC19 | 299 | 72 | 507 | 82 | 208 |
| LIF1 | 155 | 59 | 675 | 68 | 520 |
| MMM1 | 321 | 101 | 36 | 114 | -285 |
| NTA1 | 128 | 173 | 471 | 146 | 343 |
| PIN4 | 320.5 | 136 | 440 | 143 | 119.5 |
| PRY2 | 154 | 688 | 299 | 544 | 145 |
| PSP1 | 179 | 152 | 568 | 113 | 389 |
| RAD10 | 239.5 | 80 | 593 | 91 | 353.5 |
| RAD6 | 223 | 595 | 400 | 759 | 177 |
| RDI1 | 94 | 1018 | 257 | 841 | 163 |
| RPB2 | 58 | 180 | 327 | 198 | 269 |
| RSA1 | 90 | 199 | 359.5 | 194 | 269.5 |
| RTR1 | 214.5 | 112 | 669 | 89 | 454.5 |
| RTS1 | 309 | 276 | 454 | 242 | 145 |
| RVS161 | 104 | 1433 | 224 | 1292 | 120 |
| SYF2 | 62 | 134 | 248 | 193 | 186 |
| TGS1 | 161 | 191 | 393 | 287 | 232 |
| USE1 | 286 | 210 | 428 | 165 | 142 |

### Table S3. List of adjacent Intron-Containing genes localized in tandem in *S.cerevisiae*

| Upstream Gene | Downstream Gene |
|---|---|
| ARF2 | RPL35B |
| RPS9A | RPL21B |
| RPS9B | RPL21A |
| RPL19B | LSM2 |
| RPS24A | YOS1 |
| RPL22B | MOB2 |
| APE2 | RPS27A |
| RPL18B | RPS19B |
| RPL18A | RPS19A |
| YML6 | RPS18B. |

**Table S4. *S.cerevisiae* strains used in this study**

| Strain Name in Paper | Abbreviated Genotype | Full Genotype |
|---|---|---|
| Wild-Type Anchor Away | HHY168 *fpr1Δ::hphMX4* | ade 2-1 trp1-1 can1-100 leu2-3,112 his3-11,15 ura3 GAL psi+ MATalpha tor1-1 fpr1::HYG RPL13A-2×FKBP12::TRP1 (derived in house) |
| YSH1 Anchor Away | HHY168 fpr1Δ::hphMX4 YSH1-FRB::His | derived from HHY168 fpr1Δ::hphMX4 |
| RNA15 Anchor Away | HHY168 *fpr1Δ::hphMX4 RNA15-FRB::His* | derived from HHY168 fpr1Δ::hphMX4 |
| RRP6 Anchor Away | HHY168 *fpr1Δ::hphMX4 RRP6-FRB::His* | derived from HHY168 fpr1Δ::hphMX4 |
| YSH1+RRP6 Anchor Away | HHy168 fpr1Δ::hphMX4 YSH1-FRB::KanMX6 RRP6-FRB::His | derived from HHY168 fpr1Δ::hphMX4 RRP6-FRB::His |
| SNP1 Anchor Away | HHY168 *fpr1Δ::hphMX4 SNP1-FRB::kanMX6* | derived from HHY168 fpr1Δ::hphMX4 |
| Wild-Type | BY4741 | MATa his3Δ1 leu2Δ0 met15Δ0 ura3Δ0 |

**Table S5.Oligonucleotides in this study**

| Primer | Sequence (5'-3') |
|---|---|
| **RT-PCR** | |
| SEC17 RT-PCR F | AATGTCAGACCCTGTAGAGTTATT |
| SEC17 RT-PCR Int R 2.0 | GTATGTGCGCGTAAGTATATGC |
| SEC17 RT-PCR Ext R | CTACTTTGTTCTCTAAGTCATCGC |
| RPS14A RT-PCR F | CAAGAACCCGCCATGTCTA |
| RPS14A RT-PCR Int R | AATCTTCTACCTCTTCTACCACC |
| RPS14A RT-PCR Ext R | GACACGAAAAGGCGTAGAC |
| RPL18A RT-PCR F | AAGCTTACGGATTCACAATGGG |
| RPL18A RT-PCR R | AATGCGTTATTGATCGCCAG |
| RPS19A RT-PCR F Ext | AACTATGGGTTGTATATCTAATACCC |
| RPS19A RT-PCR F Int | TAAAAATGCCAGGTGTTTCCG |
| RPS19A RT-PCR R | TGTATGGTCTAACACCTCTGC |
| RPL22B RT-PCR F | CGTTTCACAAATATCAACCACA |
| RPL22B RT-PCR R | TTTATTCGTCATCCTCTTCTTCG |
| MOB2 RT-PCR F Ext | GCTTTAGCTTTTGAAACACGAAG |
| MOB2 RT-PCR F Int | TTATCAGCACATCATGTCCTTC |
| MOB2 RT-PCR R | CAAATCTATGTACTGACTAGCTGG |
| DUT1 SRB6 Fusion F | CAGATCGTTGTTGTAGACTCTCTG |
| DUT1 SRB6 Fusion R | GTTATTCACCATCATCACACTGG |
| RPL34A F | GCATAACAGAACAACCAAAGATGG |
| RPL34A Int R | TTCAGATTTCTTGGCAGCTTC |
| RPL34A EXT R | CTTCTCCAAGTCTTGGATTTCG |
| RPL26A F | GTATCAGAATGGCTAAACAATCATTAG |
| RRPL26A INT R | ATTTCGAGGAAGTTTTCAGGG |
| RPL26A EXT R | CCATTAAGGAGCTCTTCATCATC |
| ACT1 F | CCCAAGATCGAAAATTTACTGAA |
| ACT1 INT F | TTGTGGTGAACGATAGATGG |
| ACT1 EXT R | TATGATACACGGTCCAATGG |
| PDC1_ACT1 Full F | GTTGACCCAAGACAAGTCTTTC |
| PDC1_ACT1 short F | GCAAGTCAACGTTAACACCG |
| PDC1_ACT1 fusion R | CTTCATCACCAACGTAGGAGTC |

| **Color Code** |
|---|
| Figure 2 |
| Figure 3 |
| Figure 4 |
| Figure 5 |
